# An integrative model finds comparable contributions to drug resistance from poor drug adherence and adapting bacteria

**DOI:** 10.1101/773861

**Authors:** Roshan P. Mathews, Meher K. Prakash

**Affiliations:** Electrical Engineering Department, Indian Institute of Technology, Palakkad, Kerala - 678 557, India; Meher K. Prakash, Theoretical Sciences Unit, Jawaharlal Nehru Center for Advanced Scientific Research, Jakkur, Bangalore, 560 064, India

**Keywords:** Antibiotic resistance, Mutation rate, Mutation selection window, Adherence, Epidemiological model, Streptococcus pneumonia

## Abstract

Antibiotic resistance is a compound effect of several factors in the infection to healing cycle, from molecular factors such as mutation rate of bacteria to habitual behaviors such as adherence to a prescribed drug. Usually each of these factors is modeled separately from biochemistry, evolutionary biology or population health perspectives. To develop an understanding for the drug resistance at a population level, which is of high global significance, it is important to weigh all these factors in an integrated model. We develop RASAID, a model for resistance considering bacterial adaptation, infection spread, population adherence, immunity, and drug dosage. We apply the model to antibiotic resistance in the spread of resistant strains of Streptococcus Pneumoniae (*Sp*) in a finite community. We analyze the contributions from several factors to resistance, with a goal towards asking how important is the pursuit of newer drug developments relative to improving the awareness about the good practices in drug usage.

## I. INTRODUCTION

ANTIBIOTIC resistance in microorganisms is posing a threat to public health. Infection of an individual by a drug resistance is a matter of dual concern: firstly concerning their own recovery and secondly they becoming a source of transmission and spread of the resistant strain at a community level. The rise and development of resistant strains that result in longer duration of hospitalisation, higher chances of recurrent infections and mortality. While immune comprimised groups like children, elderly, diabetics etc are more sensitive, the antibiotic resistance is a global threat. At a molecular level, drug resistance in bacteria develops from bacteria developing random mutations with a certain error rate, some of which have a selection advantage against the drugs. At the population level, there are many factors that could contribute to the growth of antibiotic resistance – self medication, poor adherence to drug dose or prescription of a suboptimal dosage. Increased exposure to drugs can even arise from industrial contamination of water [1,2] and rampant antibiotic use in livestock and poultry [3].

The different pieces of the puzzle to understand the causative factors of this global crisis are spread across literature – biochemical studies interested in targeting the critical bacterial enzymes as well as identifying the key mutations in them, ecological studies comparing the selection advantage of mutants, emperical transmission-dynamic mathematical models. Over the years many attempts have been made to model antibiotic resistance at the population level [4-6] and some included the host immunity in their models [6]. Transmission-dynamic models are aimed at bridging the gap between the individual and group level effects, adherence which is a critical behavior level factor used in many clinical models is usually not factored into these models. Pharmacokinetics and bioavailability of the drug, at a dosage is usually dealt with in models which are separate from the population level models and the ones considering mutational effects. Some of the reviews [7] summarize the importance of developing mathematical models to simulate and predict various possible scenarios to arrive at strategies against drug resistance.

There is presently a surge of activity towards developing newer types of antibiotics. However, an equal emphasis is required on performing an audit of the relative contributions of the different factors to the development of resistance, failing which the resistance problem can not be comprehensively addressed. This approach requires developing integrative models which consider several aspects together, but to our knowledge there are no such models. In the present work, an integrative model for Resistance considering bacterial Adaptation, infection Spread, population Adherence, Immunity, and drug Dosage (RASAID), a schematic of which is shown in Figure 1. We apply RASAID to study the antibiotic resistance in Streptococcus pneumoniae (*Sp*) that causes contagious and acute respiratory infections. According to the World Health Organisation, pneumonia is the largest infectious cause of death among children and about 20% of the pneumonia cases in many developing countries [8]. Several strains of *Sp* are fast becoming resistant to most antibiotics and the cycle of developing newer drugs is quite long. We use RASAID to ask if the adaptation rate in bacteria is the major contributing factor and if any other anthropic origins have comparable effects on population level antibiotic resistance.

**Fig 1:**
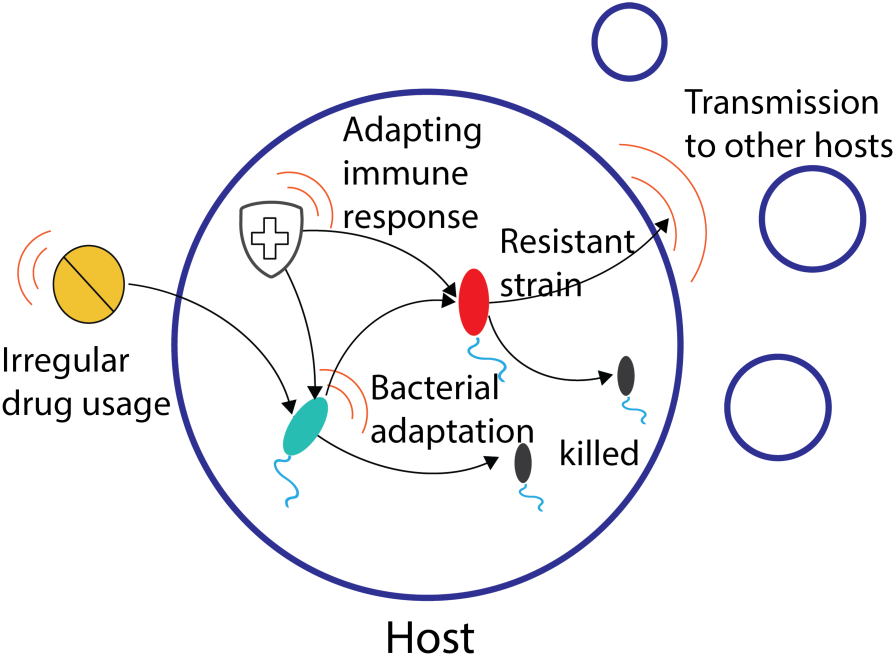
The variuos factors we consider to be contributing to the antibiotic resistance along the infection to healing process are shown in this schematic. The orange arcs represent the variability at the specific stage from infection to healing. Molecular factors such as the chance of a adaptation, evolutionary factors which give a selection advantage to the mutants, variability in drug dosage, pharmacokinetic details and adptation of immunity are also considered in a population model.

## II. INTEGRATIVE MODEL

### Major factors in the model

In this study, the development of resistant *Sp* strains was used as a subject to develop an integrative model for drug resistance. Three different strains of *Sp* are assumed based on their sensitivity relativity to cefotaxime, a commonly used antibiotic for which adequate data and relevance to usage in the public health system worldwide was found. *Sensitive* which is part of the normal nasal flora and responsive to the drug, *resistant type 1* which is an immediate strain in evolution of the bacteria challenged with cefotaxime which resists the drug but has a lower fitness because of the mutation and *resistant type 2* which maintains the resistance while regaining the fitness of the sensitive strain. The scope of the results is general and not limited to the specific choice of either the pathogen or the drug. The present model was built by adapting and integrating several available models. The logistic model for the *in vivo* (in an animal model) growth of pneumococcal bacteria [2] was updated with an adaptive immune response, where the killing of the bacteria was bacterial load dependent [6]. To estimate the bioavailability of the antibiotic, we used a first order Pharmacokinetic/Pharmacodynamic (PK/PD) elimination of the drug from the system. Poor adherence to drug was also factored into these PK/PD models and the mutant selection windows were used in the analysis based on the recommended drug dosage and usage. A hyperbolic-Monod like kill rate curve for the killing action as in [4,7] is adopted. We simulated the model for a population of 10,000 individuals. Ordinary differential equations are used to model these different populations, as well as the immune response, as detailed below. The parameters were obtained from literature as much as possible (Table 1). Where the parameters were missing, we assumed the relevant parameters that resulted in scientifically acceptable benchmarking against clinical observations such as the duration of the infection, timescales of recovery, frequency of appearance of the resistant strains etc.

**TABLE 1.** PARAMETERS AND THEIR VALUES.

| Symbol | Parameter | Value | Reference |
| --- | --- | --- | --- |
| $\alpha$ | Growth of sensitive strain determined from doubling time. | 1.386/h (30 mins)<br>(Range(20-84 mins)<br>0.495/h -2.079/h ^) | [6] |
| $k$ | Carrying capacity | $\sim 10^{12}$ | [9]^ |
| $f1$ | Fitness Factor of strain to survive antibiotic conditions (Type 1 resistance) | 0.89# | [19] |
| $f2$ | Fitness factor of compensatory mutation to restore fitness factor (Type 2 resistance) | 1.0 | [19] |
| $m1, m2$ (corresponding to $f1$ and $f2$) | Mutation rate to form resistance to bacteria | $10^{-8}$ | [15] |
| $\delta 1$ | Rate at which sensitive bacteria are killed by drug | 1.0/h | [12] |
| $\delta 2$ | Rate at which resistive bacteria are killed by drug | 0.733/h<br>(In proportion with the value used by Andreas Handel in his model [4]) | - |
| $C_{50:1}$ | Half maximal kill concentration for sensitive type | 0.25 $\mu\text{g/ml}$<br>(heterogeneous data) | [8] |
| $C_{50:2}$ | Half maximal kill concentration for all resistant type | 5 $\mu\text{g/ml}$<br>(heterogeneous data) | [8] |
| - | Mean peak concentration of antibiotic in otitis media | 5 $\mu\text{g/ml}$ | [11] |
| $\lambda$ | Half life of Cefotaxime | 1.1h | [20] |
| $s$ | Saturation point of immune kill rate | $k/100$ | [8] |
| $x$ | A Gaussian factor for rate at which the immunity kills pathogen to bring in variation in a population. | $mean=1, variance=0.06$ | @ |
| $d$ | Immune response decay at a fixed rate | 20** | @ |
| $g$ | a) Healthy<br>b) Incubation<br>c) First Drug dosing period<br>d) Subsequent drug dosing period<br>e) Infected state | a) $\alpha * 10^5$<br>b) $\alpha * 0.2 * 10^3$<br>c) linear increase from $\alpha * 0.2 * 10^3$ to $\alpha * 10^4$ in 168 hours<br>d) $\alpha * 10^4$<br>e) $\alpha * 10^4$ | @ |
| $b$ | Rate constant at which immunity kills bacteria (Sp)<br>a) Healthy<br>b) Incubation<br>c) First Drug dosing period (L)<br>d) Subsequent drug dosing period<br>e) Infected state | a) $10^*s$<br>b) $s$<br>c) $s$<br>d) $s$<br>e) $s$ | @ |

^Data based on mouse used to model in vivo condition
\*\*The decay of the immune response is assumed to be fast that it will closely follow the bacteria load
#Range for in-vivo fitness of type 1 resistance is 0.7-0.89. To pronounce the effect of its presence in the simulation we have considered the upper limit. The lower fitness factors don't give suitable simulation results for arriving to conclusions. It is probably this reason that the menace of antibiotic resistant strains is constrained to smaller numbers in the society and not leading to a mass outbreak. The higher fitness factor leads to larger generation of resistant strains and can be used to figuratively and logically predict trends.

### Bacterial growth

Since *Sp* is a natural flora of the respiratory tract, from studies [6,9] we have adopted a normal count as about 10^6^ sensitive strains and 0 resistant strains at the start of the simulation. A person is considered infected if the bacterial numbers rise by a 100x of the normal level, which corresponds to the numbers found in mouse models [9,10]. At the level of the individual, the growth of the bacteria was assumed to be according to a logistic growth [2] as in [Eq1] below.

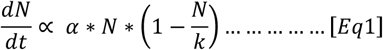

where *N* is the number of bacteria, *α* the growth rate and *k* is the carrying capacity. Extrapolating the numbers found in mice to humans, a carrying capacity of 10^12^ was assumed (justification in Supporting information).

### Drug dosage, mutation selection

The typical therapeutic dosage of Cefotaxime for an adult is about 500 to 1000mg, twice or thrice a day depending on the medical condition of the patient. We assume the drug is given for 7 days (2 times a day) with an extension period of 4 days if required (maintaining the same dosage as before) and extension period up to three extensions in a month. From the experimental study conducted [11] the peak antibiotic concentration in otitis media exudates ranges from 2-10 µg/ml for therapeutic doses.

In the current study, we assumed that the resistance development is correlated with the higher minimum inhibitory concentration (MIC). The mutations are taken in dual stages; the first being the mutation in PBP2x for the immediate resistance to the drug and a compensatory mutation in PBP1a after the drug pressure is removed. Although the mutations mentioned above are not necessarily point mutations, for the sake of simplicity in modeling it, we have considered them to be point mutations.

### Antibiotic action

The drug concentration *in vivo* is modeled by using the first order pharmacokinetics [Eq2]

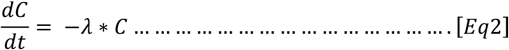

where, *λ* is the half-life of the drug. The rate at which the drug kills the sensitive and resistant bacteria are sigmoidal as below [5,6].

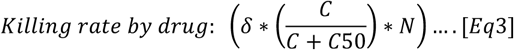

In equation [Eq3], δ is the rate at which the drug kills sensitive bacteria; *N* is the susceptible bacteria population. *C*_*50*_ is the concentration at which half the maximum kill is achieved.

The maximal killing rate of sensitive bacteria is taken to be 1 as in [12] and the maximal killing rate of resistant bacteria has been assumed to be in proportion to the value adopted in [8], parametrizing it with the maximal killing rate from the above mentioned study on cefotaxime [12]. The rest of the parameters are kept the same as in the model described in [13]. It has been reported in the literature that the mutant selection occurs in cephalosporins at concentrations less than or equal to 16 times MIC [13,14]. This level is termed as MPC (mutant prevention concentration) above which chance of resistance buildup is negligible. For single step mutations the MPC for Cefotaxime was determined to be about ∼ 8*MIC [15]. For the above MPC, to gain resistance to antibiotics multiple mutations have to occur thus decreasing the chance of resistance buildup. For concentrations below MIC, the drug selection pressure is not sufficient to cause mutation [18]. *Sp* sensitive to cefotaxime are defined as a colony with the MIC <= 0.25 µg/ml [16]. The effect of this mutant selection window has been modeled with a higher mutation rate with the concentration coming within the mutant selection window and a lower mutation rate when it falls outside of it.

### Adaptive immune response

We have adopted the model described by [Eq2] where the immune response is both the function of bacterial load dependent immune response dynamics and bacterial load dependent immune response killing.

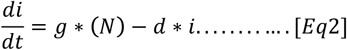

where *g* and *d* are constants.

Killing term of pathogenic bacteria by immunity:

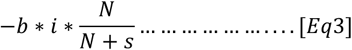

where *i* represents the immune response term, N is the total amount of bacteria present, *s* is the saturation of the immune response and *b* is the rate constant at which the bacteria are eliminated by the immune system. The values of the constants used in the equation are given in Table 1.

Some of the parameters used are estimated as follows. In this work, the basic dictum in benchmarking the immunity level is that in a healthy person the immune response is such that it keeps the bacterial growth in check. The incubation for *Sp* when the symptoms are clearer and the patient is recommended an antibiotic dosage generally varies between a few hours to 2 days, and we use the latter in our model. Generally, a reduction in immunity could be an aiding circumstance for getting an infection. For simulating this aspect, we decrease the constants in the immunity by a few factors of 10’s. For the recovery of immunity, we initially increase it in a linear fashion during the first dosage of the drug to a level where the immunity manages to keep the bacterial load just on the border line of infection threshold (such that there is a necessity of drug action to heal the patient) and for the subsequent time we increase them in steps. A random Gaussian variable with a mean of 1 and variance of 0.06 was introduced to bring in a sense of variation among the population and to make the model more realistic by multiplying this factor with our estimated parameter for immunity.

### Infection spread

The infection is assumed to spread from the individual in the following way. A *contact rate* which describes the number of contacts made by an individual in a particular duration. Although this contact rate could be variable [17], an approximate rate of 40 contacts per month, which can potentially transfer bacteria from to another was assumed. At the end of each month in our simulation, we select 40 members at random from the population with whom an individual makes contact (handshakes). The number of bacteria transferred are proportional to the number of bacteria the individual has [18], thus bringing an asymmetry to the transmission between individuals of different infection levels.

### Overall model at the population level

We combine all the factors discussed above into a population level model for 10,000 individuals. An average of eighty new cases per month was assumed. The set of simultaneous ordinary differential equations (ODEs) was developed and simulated for a period of 12 months.

#### a) Healthy state, Incubation state and Infected state

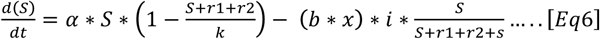

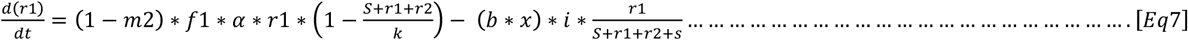

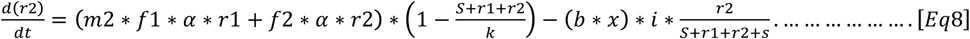

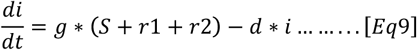

#### b) During drug intake duration

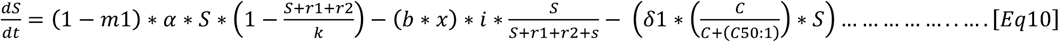

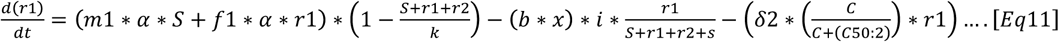

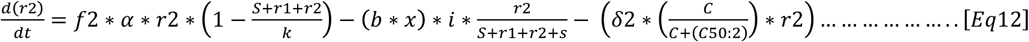

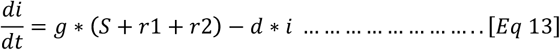

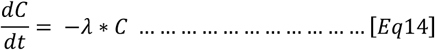

In the above equations, the notation, *S*-sensitive, *r1*-resistance formed during antibiotic environment, *r2*-compensatory mutation was used. It may be noted that for healthy state, incubation state and infected state, the structure of the system of equations remain unaltered, however the numerical values are subject to the stated considerations as mentioned in Table 1. The system of simultaneous equations was solved numerically using Python.

## III. RESULTS AND DISCUSSIONS

### A. Growth and Spread of Resistant strain (within host)

We simulated the numbers of bacteria of the three types. The fitness of resistant types 1 and 2 was assumed to be 0.89 and 1 respectively. The simulation was performed for a time period of one month and the results for these bacterial numbers assuming drug usage adherences from 60% to 100% are plotted in Figure 2.

**Fig 2:**
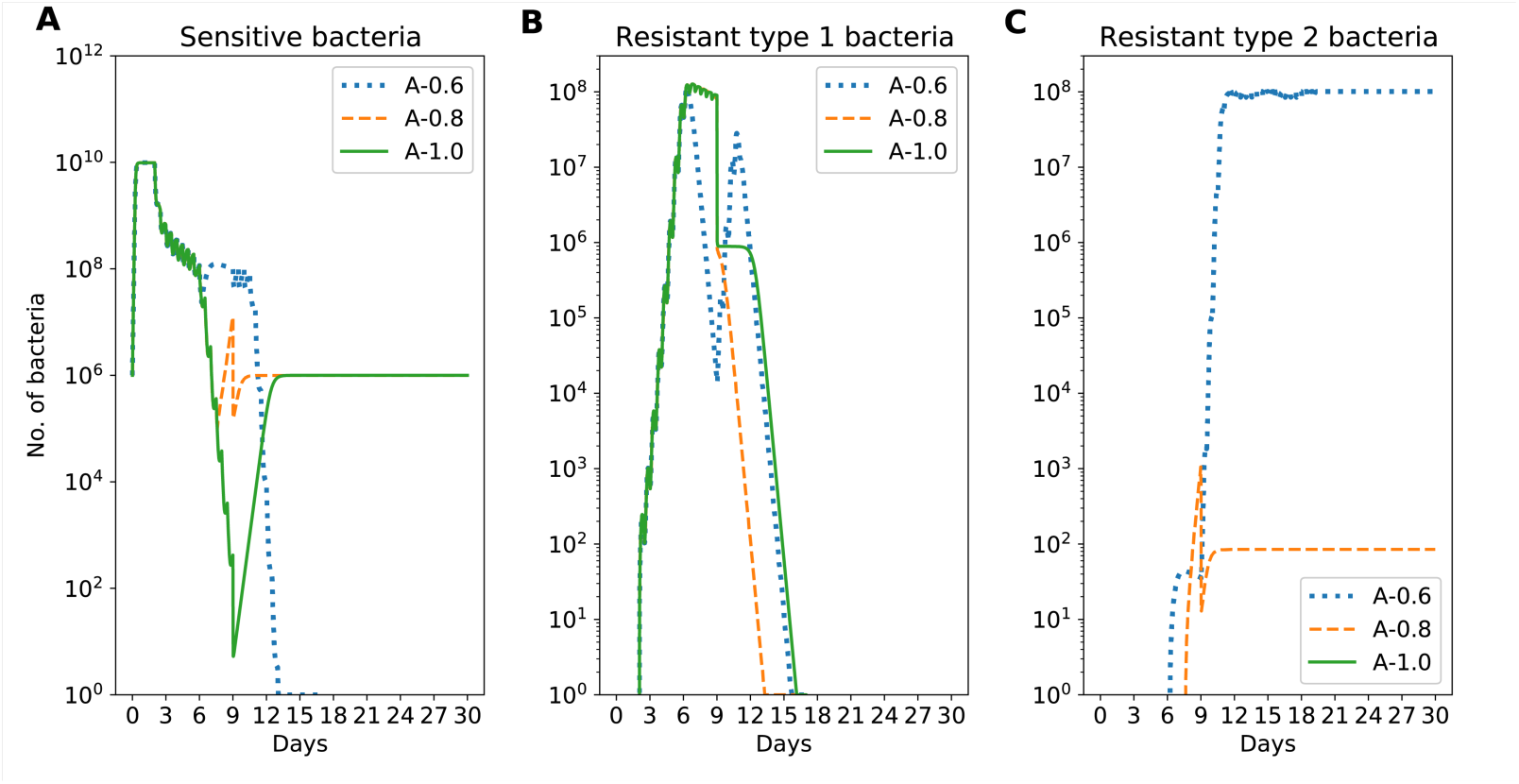
Studying the growth of bacteria (sensitive & resistant) in an individual for different adherence levels. The simulation starts with base level bacterial load of 10^6^ in the individual. The Gaussian factor (x) for the immunity was taken to be 1 for this simulation. The calculations were repeated for three different cases where the individual adheres to 60%, 80% or 100% (A0.6, A0.8, A1.0 respectively) of the prescribed dosage. At the recommended dosage for a normal healthy person, there is no resistance type 2 strain at the end of the infection period. However, as it can be seen in (C), when the adherence is poor, it leads to the development of resistant type 2.

In Figure 2, panel A shows the number of sensitive bacteria in the individual throughout the period of a month. As seen in the plot, in the initial stage of incubation (48 hours), the individual maintains a high level of sensitive strains and upon the antibiotic pressure, the count of the sensitive bacteria fall drastically responding to the treatment. In perfect adherence (equal to 1, represented by solid line) sensitive strains falls to the minimum value under antibiotic pressure towards the end of the therapeutic period. Due to its higher multiplication rate when compared to the resistant strains, it regains itself to the benign count level of the default nasal bacterial flora (growth is balanced by immune response). In partial adherence (of 0.8), as the antibiotic pressure eases out earlier, the number of sensitive strains follows the trend of perfect adherence over a period of time however has a sudden increase due to the relieving of the antibiotic pressure (Fig. 3B). However in the case of adherence of 0.6, we notice a different trend contrary to the common expectation. Here the number of sensitive strains falls to a negligible level even though the antibiotic is discontinued at early stage of the treatment. This is due to the repeated exposure (extension periods) and with the growth of resistant strains. As shown in panel B, in perfect adherence, the number of resistant type 1 progressively builds up as long as the antibiotic pressure exists and subsequently drops to negligible level in about 16 days due to the low fitness factor. The rate at which the resistant strain reproduces is much slower than the rate at which immunity kills them and the competition for resources from the growing sensitive strain depicted well in (Figure 3) is responsible for their eradication. In the cases of partial adherence, due to their imperfect adherence to the first duration of treatment the individual remains infected and takes an additional dosage of drug namely the extension period mentioned in the materials and methods. Due to this a secondary spike is seen in the number of resistant type 1 bacteria in the individual. Due to imperfect adherence causing recurrent infections the immune system is weakened and the antibiotic exposure time is increased. Decrease in the antibiotic concentration in the individual during this time due to poor adherence gives rise to favorable conditions for compensatory mutations to develop restoring the fitness cost imposed by the drug resistance.

**Fig 3:**
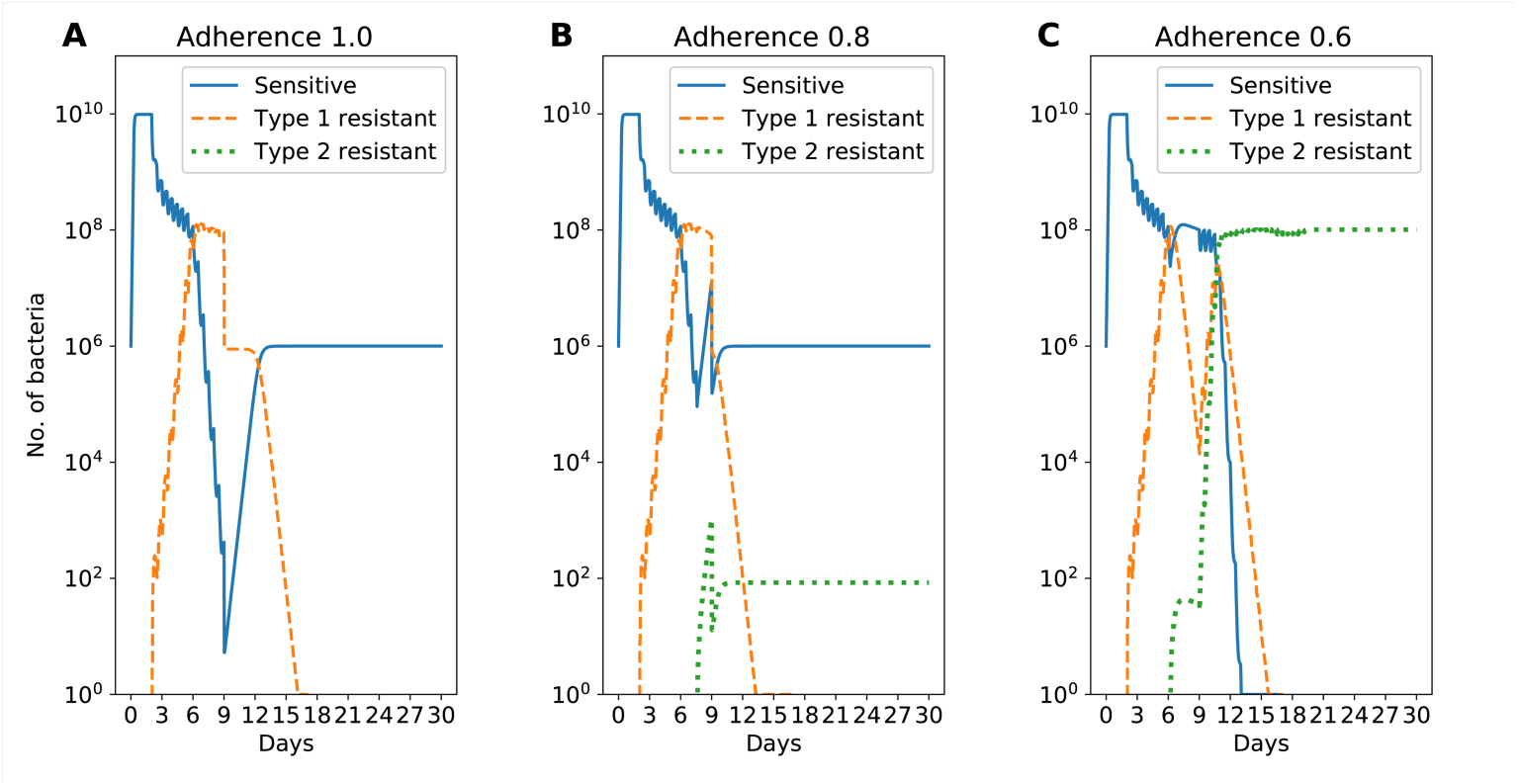
The data is exactly same as in Fig 2. However, it is presented from a different perspective. All the data corresponding to one adherence are pooled together in each of the subplots.

As seen in panel C, with the perfect adherence of 1, the number of resistant strains developed is not in great numbers and the immune system comes back to normal by then. This as seen from the study by [8] reduced the mutant selection window and thus very few resistant type 2 strains are observed in the simulation. Whereas in the case of lower adherences such as 0.8 and 0.6, as explained above the favorable conditions like a still recovering immunity broadens the mutant selection window by the slower killing of the bacteria thus increasing the probability of the mutation occurring. This is also exacerbated by the lack of competition from the sensitive strains that are significantly depleted from the drug dosage and would take time to recover to their benign level. Thus, we are able to see the stark increase in numbers of resistant type 2 strains in the population for imperfect adherences. All these are under the assumption that seven days of the drug are required for cure. For this particular immunity level chosen, seven days are required. In the case of a person with a lower immunity level (immuno-compromised) seven days may be a non optimal dose.

### B. Growth and Spread of Resistant strain in a Population with fixed adherence

The simulation was then repeated for a population of 10,000 people. The immunity distribution in the population was assumed to be a Gaussian (Table 1). From panel B, we can quantitatively see that the number of resistant type 2 bacteria in the population is lesser in number with good adherence. The reason for the sudden spurt of the resistant bacteria might be the possibility of an immune-compromised person getting infected. In case of an immune-compromised person as studied in [13], the resistant strains grow uninhibited by the lack of competition from sensitive strains as they get killed by the drug and a weak immunity aids the proliferation of resistant strains. Panel C, shows the sensitive bacteria in the population. With complete adherence the sensitive strain is the highest (compared to imperfect adherence) resulting in the resistant bacteria to be minimum due to the competition faced. This also is a better scenario as the sensitive bacteria being in the majority in the population makes the treatment effective in case of an illness. The higher the number of resistant strains in the population, the higher the chances of the population being infected by the resistant strain and this case is of great public health concern as the drugs present in the market have lesser effect on them. Panel D is the overall indicator of the effectiveness of treatment in the population. Panel D, shows the number of people infected in each month. In the case where all infected people are completely healed the plot would be a straight line at 8 cases of infection per month. The line deviating from the normal level shows us that there are people infected at the end of the month and aren’t fully cured. In the event that the person cannot be cured in the same month he is carried forward to the next month and continued.

### C. Growth and Spread of Resistant strain in a Population with mixed adherence

In a population, a more likely scenario than a pure adherence of 0.6 or 0.8 is a distribution of individuals with different adherences. We assumed three different distributions (Figure 5) and studied the dependence of the development of resistant strains as well as the number of infected individuals. It is clear that the communities which have a better distribution of adherence are more effective in containing the infections.

**Fig 4:**
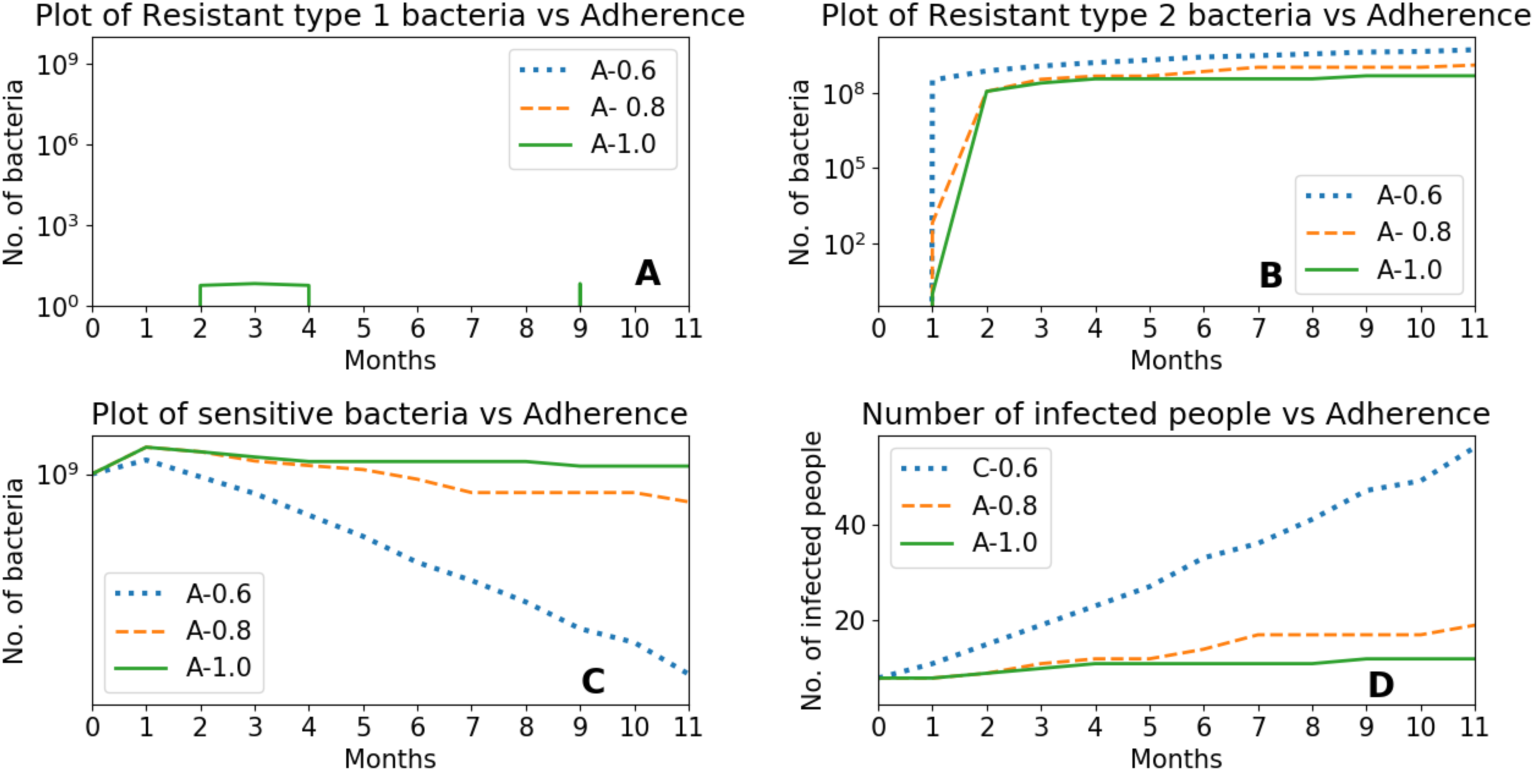
The bacteria growth and dynamics for varying adherences in a population. Panels A to C represent the average number of bacteria in the population. Panel D is the total number of infected individuals. The adherence has a strong influence at the population level.

**Fig 5:**
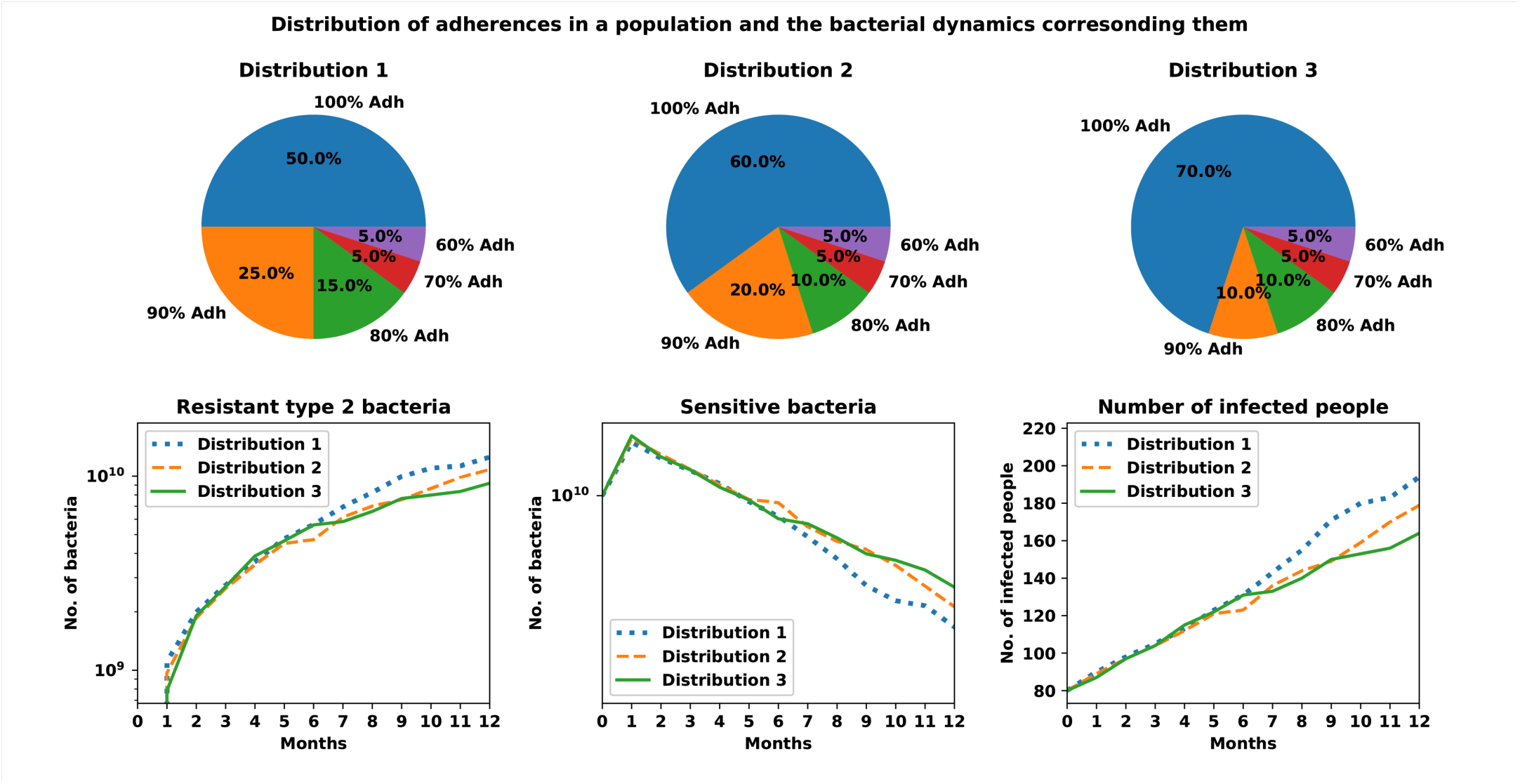
Distribution (hypothetical) of adherences in a population was assumed as shown in the pie chart. The evolution of the community level resistance was studied assuming these distributions. More compliant communities have better chances of fighting the antibiotic resistance.

### D. Comparing the effect of variation in immunity and adherence

In the calculations we performed at the population level, we sorted the number of people infected by resistant strain type 2 according to their innate immunity and the adherence. It is clear from Fig 6 that the it is important for the immune compromised to have better adherence. The results also suggest that a moderate reduction in immunity can be compensated by a better adherence to the optimal dosage.

**Fig. 6.**
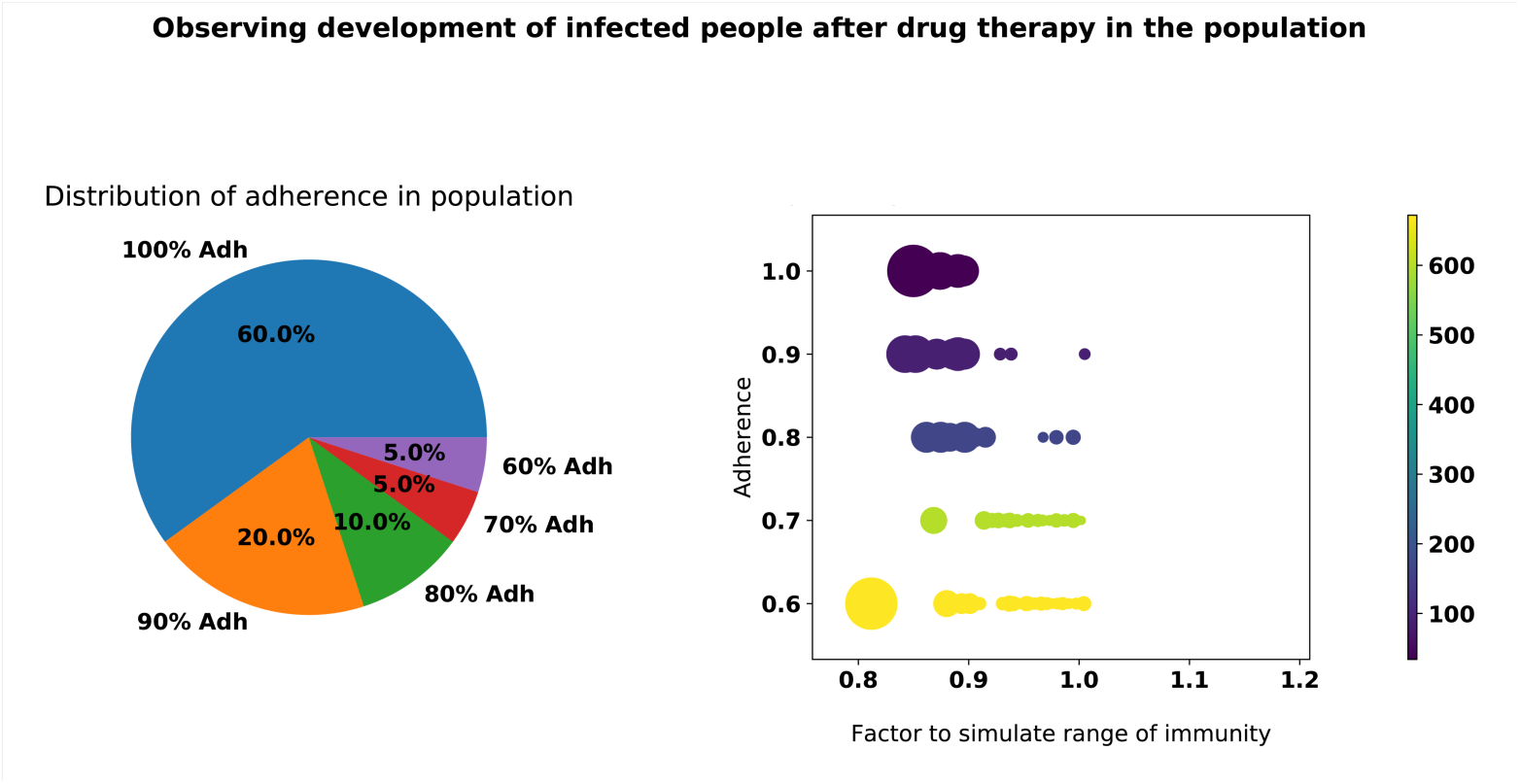
For a population with mixed adherence shown on the left, the number of people infected with resistant strain 2 was calculated depending on the adherence and innate immunity levels. The color bar indicates the number of infected. It is clear from the graph that a moderate reduction in immunity can be compensated by a strict adherence.

### E. Comparing the contributions from bacterial adaptation with adherence

We also compared the rise in number of infected people (Figure 7) when the adherence was perfect, but mutation rates increased by a factor of 100, to those when the adherence dropped to 0.6. Interestingly, while the effects of an adherence of 0.8 was comparable to a 100x increase in the mutation rates, adherence of 0.6 significantly increases the number of infections, from the resistant strains. The finding underscores the need to comprehensively evaluate all the possible factors that contribute to the antibiotic resistance.

**Fig 6:**
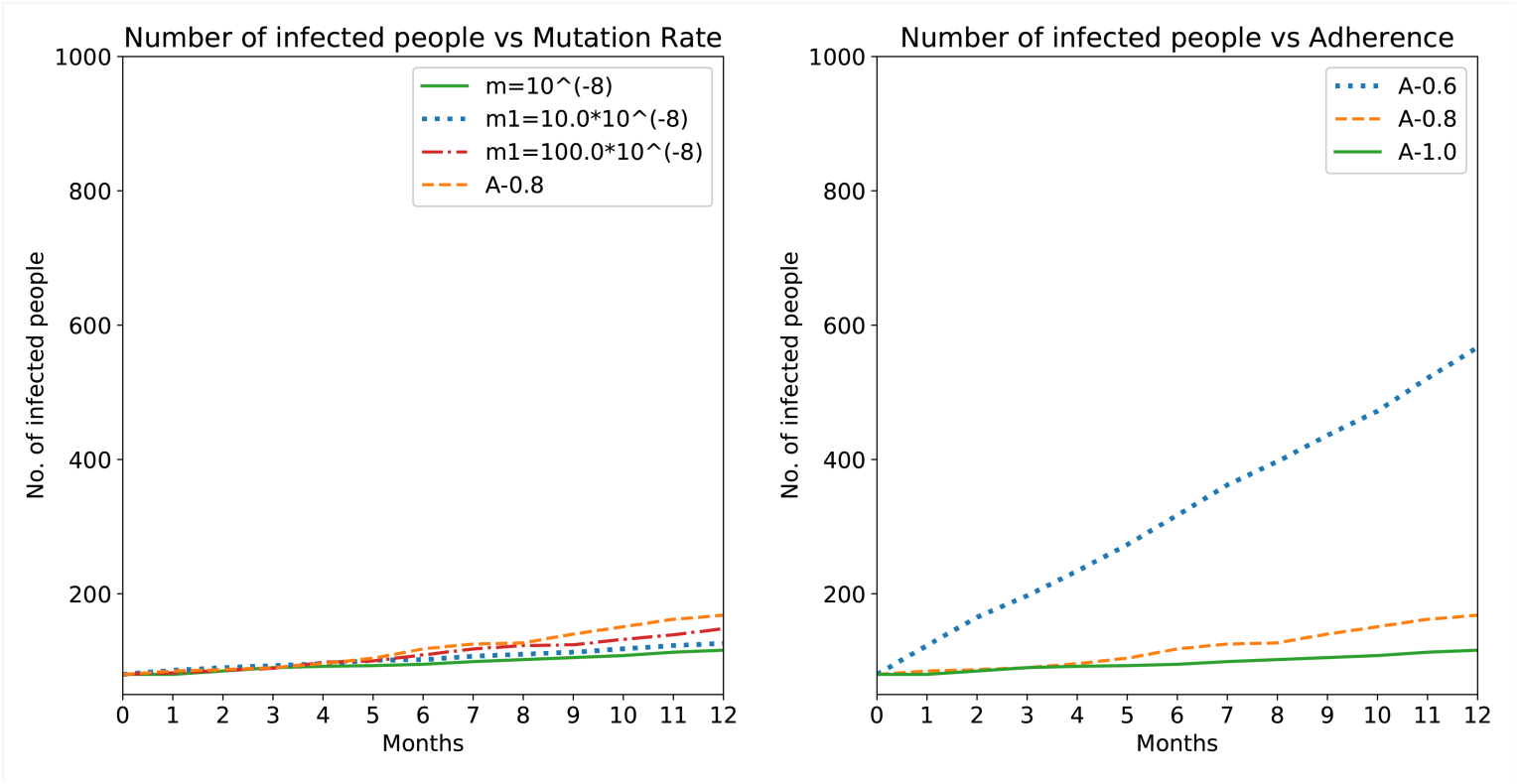
Comparison of the number of infected people when (A) mutation rates increased significantly (B) the adherence dropped to 0.6. In (A), the results from A=0.8 is given for a reference.

The limitations of the model are that it assumes that the treatment course continues to be the same and no new antibiotics are suggested to a patient that is not responding. While the model studies the competition between the different strains, we do not consider competition from any other bacterial types not related to *Sp*. Further, the since we studied *Sp*, one expects to have seasonal variations, which is not taken into account in the present model. Despite these limitations, we believe the integrative model we present combines information across scales from mutations, mutant selection window, pharmacokinetics, drug dosage and adherence to quantify the different contributions to the population level antibiotic resistance.

## IV. CONCLUSION

We developed an integrative model to scan across the contributors from bacterial adaptation rate to drug dosage adherence by patients. The model allowed us to weigh the factors contributing to the resistance development. The comprehensive model allowed us to see that adherence is comparable, if not more important to the mutational adaptation of bacteria. Considering the urgency of tackling the antibiotic resistance, we believe the integrative model gives an opportunity to weigh the cost-benefits of spending resources on the different factors contributing to the drug resistance.

## V. ACKNOWLEDGEMENTS

RPM acknowledges the Summer Research Fellowship Program of the Jawaharlal Nehru Centre for Advanced Scientific Research, which allowed performing the present research.

## VI. SUPPORTING INFORMATION

The Python scripts used for the analysis will be available upon request.

## REFERENCES

[1] Fick, J., Söderström, H., Lindberg, R. H., Phan, C., Tysklind, M., & Larsson, D. J. (2009). Contamination of surface, ground, and drinking water from pharmaceutical production. Environmental Toxicology and Chemistry, 28(12), 2522–2527.

[2] Davies, J., & Davies, D. (2010). Origins and evolution of antibiotic resistance. Microbiol. Mol. Biol. Rev., 74(3), 417–433.

[3] Landers, T. F., Cohen, B., Wittum, T. E., & Larson, E. L. (2012). A review of antibiotic use in food animals: perspective, policy, and potential. Public health reports, 127(1), 4–22.

[4] Smith, A. M., McCullers, J. A., & Adler, F. R. (2011). Mathematical model of a three-stage innate immune response to a pneumococcal lung infection. Journal of theoretical biology, 276(1), 106–116.

[5] Regoes, R. R., Wiuff, C., Zappala, R. M., Garner, K. N., Baquero, F., & Levin, B. R. (2004). Pharmacodynamic functions: a multiparameter approach to the design of antibiotic treatment regimens. Antimicrobial agents and chemotherapy, 48(10), 3670–3676.

[6] Handel, A., Margolis, E., & Levin, B. R. (2009). Exploring the role of the immune response in preventing antibiotic resistance. Journal of theoretical biology, 256(4), 655–662.

[7] Andersson, D. I., & Hughes, D. (2011). Persistence of antibiotic resistance in bacterial populations. FEMS microbiology reviews, 35(5), 901–911.

[8] World health organization: Pneumonia factsheet: https://www.who.int/news-room/fact-sheets/detail/pneumonia (accessed on 21st June, 2019)

[9] Francis, K. P., Yu, J., Bellinger-Kawahara, C., Joh, D., Hawkinson, M. J., Xiao, G., … Contag, P. R. (2001). Visualizing Pneumococcal Infections in the Lungs of Live Mice Using Bioluminescent Streptococcus pneumoniae Transformed with a Novel Gram-Positive luxTransposon. Infection and immunity, 69(5), 3350–3358.

[10] Smith, A. M., Adler, F. R., Ribeiro, R. M., Gutenkunst, R. N., McAuley, J. L., McCullers, J. A., & Perelson, A. S. (2013). Kinetics of coinfection with influenza A virus and Streptococcus pneumoniae. PLoS pathogens, 9(3), e1003238.

[11] Novick Jr, W. J. (1982). Levels of cefotaxime in body fluids and tissues: a review. Reviews of infectious diseases, 4 (Supplement_2), S346–S353.

[12] Nix, D. E., Wilton, J. H., Hyatt, J., Thomas, J., Strenkoski-Nix, L. C., Forrest, A., & Schentag, J. J. (1997). Pharmacodynamic modeling of the in vivo interaction between cefotaxime and ofloxacin by using serum ultrafiltrate inhibitory titers. Antimicrobial agents and chemotherapy, 41(5), 1108–1114.

[13] Canet, J. J., & Garau, J. (2002). Importance of dose and duration of β-lactam therapy in nasopharyngeal colonization with resistant pneumococci. Journal of Antimicrobial Chemotherapy, 50(Suppl_3), 39–44.

[14] Drusano, G. L., & Goldstein, F. W. (1996). Relevance of the Alexander Project: pharmacodynamic considerations. Journal of Antimicrobial Chemotherapy, 38(Suppl_A), 141–154.

[15] Sifaoui, F., Kitzis, M. D., & Gutmann, L. (1996). In vitro selection of one-step mutants of Streptococcus pneumoniae resistant to different oral beta-lactam antibiotics is associated with alterations of PBP2x. Antimicrobial agents and chemotherapy, 40(1), 152–156.

[16] Jacobs, R. F., Kaplan, S. L., Schutze, G. E., Dajani, A. S., Leggiadro, R. J., Rim, C. S., & Puri, S. K. (1996). Relationship of MICs to efficacy of cefotaxime in treatment of Streptococcus pneumoniae infections. Antimicrobial agents and chemotherapy, 40(4), 895–898.

[17] Blackwood, J. C., & Childs, L. M. (2018). An introduction to compartmental modeling for the budding infectious disease modeler. Letters in Biomathematics, 5(1), 195–221.

[18] Lingaas, E., & Fagernes, M. (2009). Development of a method to measure bacterial transfer from hands. Journal of hospital infection, 72(1), 43–49.

[19] Orio, A. G. A., Piñas, G. E., Cortes, P. R., Cian, M. B., & Echenique, J. (2011). Compensatory evolution of pbp mutations restores the fitness cost imposed by β-lactam resistance in Streptococcus pneumoniae. PLoS pathogens, 7(2), e1002000.

[20] Fu, K. P., Aswapokee, P. R. A. S. I. T., Ho, I. R. W. I. N., Matthijssen, C. H. A. R. L. E. S., & Neu, H. C. (1979). Pharmacokinetics of cefotaxime. Antimicrobial agents and chemotherapy, 16(5), 592–597.

